# Comprehensive Assessment of Somatic Copy Number Variation Calling Using Next-Generation Sequencing Data

**DOI:** 10.1101/2021.02.18.431906

**Authors:** Yun-Ching Chen, Fayaz Seifuddin, Cu Nguyen, Zhaowei Yang, Wanqiu Chen, Chunhua Yan, Qingrong Chen, Charles Wang, Wenming Xiao, Mehdi Pirooznia, Daoud Meerzaman, The Somatic Mutation Working Group of the SEQC-II Consortium

## Abstract

Copy number variation (CNV) is a common type of mutation that often drives cancer progression. With advances in next-generation sequencing (NGS), CNVs can be detected in a detailed manner via newly developed computational tools but quality of such CNV calls has not been carefully evaluated. We analyzed CNV calls reported by 6 cutting-edge callers for 91 samples which were derived from the same cancer cell line, prepared and sequenced by varying the following factors: type of tissue sample (Fresh vs. Formalin Fixed Paraffin Embedded (FFPE)), library DNA amount, tumor purity, sequencing platform (Whole-Genome Sequencing (WGS) versus Whole-Exome Sequencing (WES)), and sequencing coverage. We found that callers greatly determined the pattern of CNV calls. Calling quality was drastically impaired by low purity (<50%) and became variable when WES, FFPE, and medium purity (50%-75%) were applied. Effects of low DNA amount and low coverage were relatively minor. Our analysis demonstrates the limitation of benchmarking somatic CNV callers when the real ground truth is not available. Our comprehensive analysis has further characterized each caller with respect to confounding factors and examined the consistency of CNV calls, thereby providing guidelines for conducting somatic CNV analysis.

## Introduction

Cancer in many ways is a disease of the genome. It evolves with accumulations of various types of mutations and can have a genetic and epigenetic background of heritable factors [1–3]. Tumor-derived CNVs are among the most important genomic aberrations in cancers together with somatic mutations and translocation/rearrangement structural variants (SVs). Since a CNV amplification is often associated with oncogene activation, and a CNV deletion is frequently attributed to the gene inactivation [4–6], an accurate detection of CNVs plays a crucial role in cancer diagnosis and treatment.

Several technologies such as Fluorescence In Situ Hybridization (FISH) [7], NanoString’s digital detection technology [8], arrayCGH [9], and SNP array [10] have been widely used in detecting somatic CNVs. Recently, Next Generation Sequencing (NGS)-based platforms have been emerging as a primary tool to detect CNVs [11–13]. NGS can potentially provide a detailed picture of the CNVs that characterize the tumor. Copy number can be inferred from NGS data by calculating the depth ratio of a given genomic position compared to the rest of the genome or a control sample [14].

In recent years, many bioinformatics tools have been developed for somatic CNV calling based on NGS data but the quality of their CNV calls has not been carefully evaluated yet. Calling CNVs for cancer data is a challenging task. It can be complicated by many confounding factors such as the amount and type of input DNA for library preparations, presence of normal cells in the tumor specimen (tumor purity), intra-tumor heterogeneity, and sequencing platform and coverage [15]. A few studies have performed method evaluation for CNV callers. However, they focus on either non-cancer data [16] or a specific data type (whole-exome sequencing data) [17, 18] and simply benchmark callers using conventional metrics (such as specificity and sensitivity) without investigating how potential confounding factors would affect somatic CNV calling. Given the high level of heterogeneity in cancer data, a comprehensive assessment is crucial to better understand strength and weakness of state-of-art somatic CNV callers.

To this end, we evaluated 6 cutting-edge somatic CNV callers – ASCAT [19], CNVkit [20], FACETS [21], Control-FREEC [22], Segmentum [23], and Sequenza [14] – using a collection of 91 samples derived from the same tumor-normal paired cancer cell line, in which replicates were prepared and sequenced by varying the following factors: the amount of input DNA for library preparation, type of the tissue (Fresh vs. Formalin Fixed Paraffin Embedded (FFPE)), tumor purity, sequencing platform (Whole-Genome Sequencing (WGS) versus Whole-Exome Sequencing (WES)), and sequencing coverage. With this dataset, we were able to benchmark callers’ performance, evaluate consistency of CNV calling across replicates, and characterize each caller by interrogating alterations of CNV calls regarding each confounding factor. With this comprehensive analysis, we hope to provide guidance for discovering somatic CNVs using NGS-based CNV calling and future directions for improving CNV detection methods.

## Results

### Sample preparation and analysis workflow

We used the SEQC-II reference matched samples, a human triple-negative breast cancer cell line (HCC1395) 291 and a matched B lymphocyte derived normal cell line (HCC1395BL) in our analysis. Detailed sample information can be found in the SEQC-II reference samples manuscripts [24, 25]. We used TruSeq library for 21 sample preps with optimum amount of input DNA (1000ng fresh tumor DNA with 100% tumor purity) and performed WGS across different sequencing sites. This optimum set was used as the “reference” (REF) to benchmark the performance of CNV callers and to assess the effect of the confounding factors. Another set of data comprised replicates affected by varying each of the following confounding factors: input DNA amount, FFPE-processed sample (versus fresh samples), tumor purity, sequencing coverage, and WES (versus WGS). A total of 91 samples were prepared as summarized in Figure 1A. In addition, we also interrogated the effect of computational methods by calling CNVs for these 91 samples using 6 somatic CNV callers: ASCAT, CNVkit, FACETS, Control-FREEC (abbreviated as FREEC), Segmentum, and Sequenza, totaling 546 (= 91*6) sets of genome-wide CNV calls. These CNV calls were analyzed as follows (Figure 1B): (1) testing consistency of CNV calls among 21 REF replicates within and between callers as well as sequencing sites; (2) comparing CNV calls of confounded samples to those of REF samples to assess effects of confounding factors; and (3) benchmarking performance of CNV callers based on the 2-out-of-3 majority rule, the array-based CNV calls, and a set of high confidence CNV calls. To compare CNV calls across callers, we divided the genome into 287,509 bins with bin size of 10KB and assigned each bin a copy number (CN) based on CNVs reported by each caller for each sample (see Methods). In addition to actual CNs, we also categorized each bin into amplification (CN > 2.5), deletion (CN < 1.5), or marginal (1.5 <= CN <= 2.5). CNs of 287,509 bins for 548 sets of CNV calls (546 sets of NGS calls + 2 sets of array calls) were archived in Supplementary Table 1.

**Figure 1.**
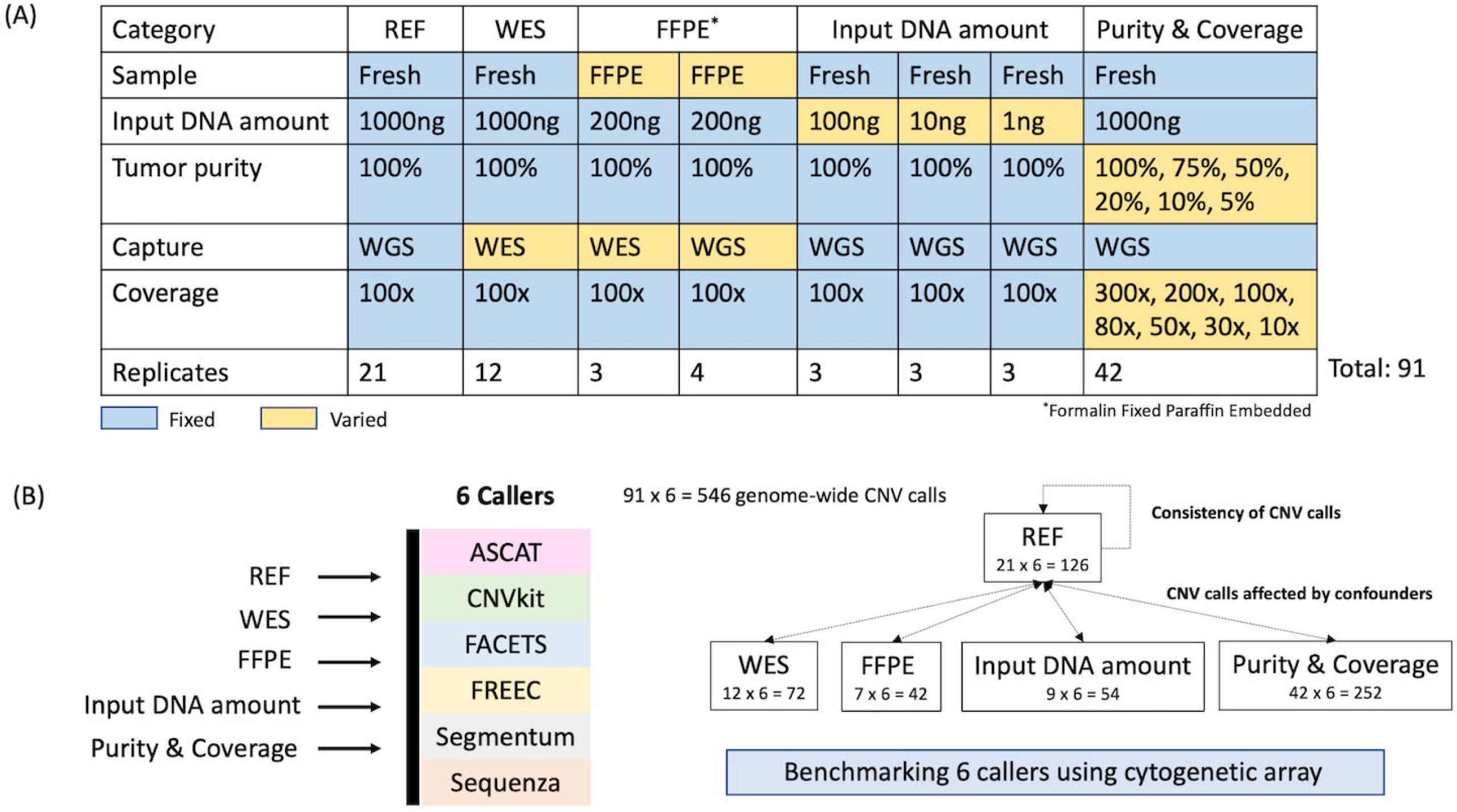
(A) Sample preparation and (B) analysis workflow.

### Between-caller variation of CNV calls is mainly driven by CNV callers but not sequencing sites whereas within-caller variation coincided with highly amplified regions

We started with the 21 REF samples to evaluate the reproducibility within and between callers as well as sequencing sites without any effects of confounding factors. We categorized the median CN of each bin (across the 21 REF samples) into marginal, amplification, and deletion, and observed that the proportion of amplifications and deletions detected by ASCAT, FACETS, FREEC, and Sequenza follow a similar pattern with more amplifications identified, while more deletions were detected by CNVkit and Segmentum (Figure 2A). This shows the overall disagreement in calling amplification and deletion between callers. The principle component analysis (PCA) on CNs indicated that variation among CNV calls was mainly driven by callers but not sequencing sites (Figure 2B). However, pairwise distances between different sites in the PC space were significantly larger than those within the same sites (One-sided Wilcoxon rank sum test p<0.05 for all callers) (Figure 2C) for all callers, suggesting sequencing sites still contributed to variation of CNV calls though relatively minor compared to callers. Despite the disagreement in calling amplification and deletion between CNVkit and Segmentum versus the other 4 callers, their median CNs were highly correlated (Spearman correlation) across the whole genome between all callers (upper triangular matrix in Figure 2D). This suggests that the trend of CNs across bins was consistent between all 6 callers (i.e. bins with higher CNs in one caller were also with higher CNs in another caller relative to other bins) but the absolute CNs were called lower in CNVkit and Segmentum yielding more bins with CNs less than 2 and, thus, categorized as deletions.

**Figure 2.**
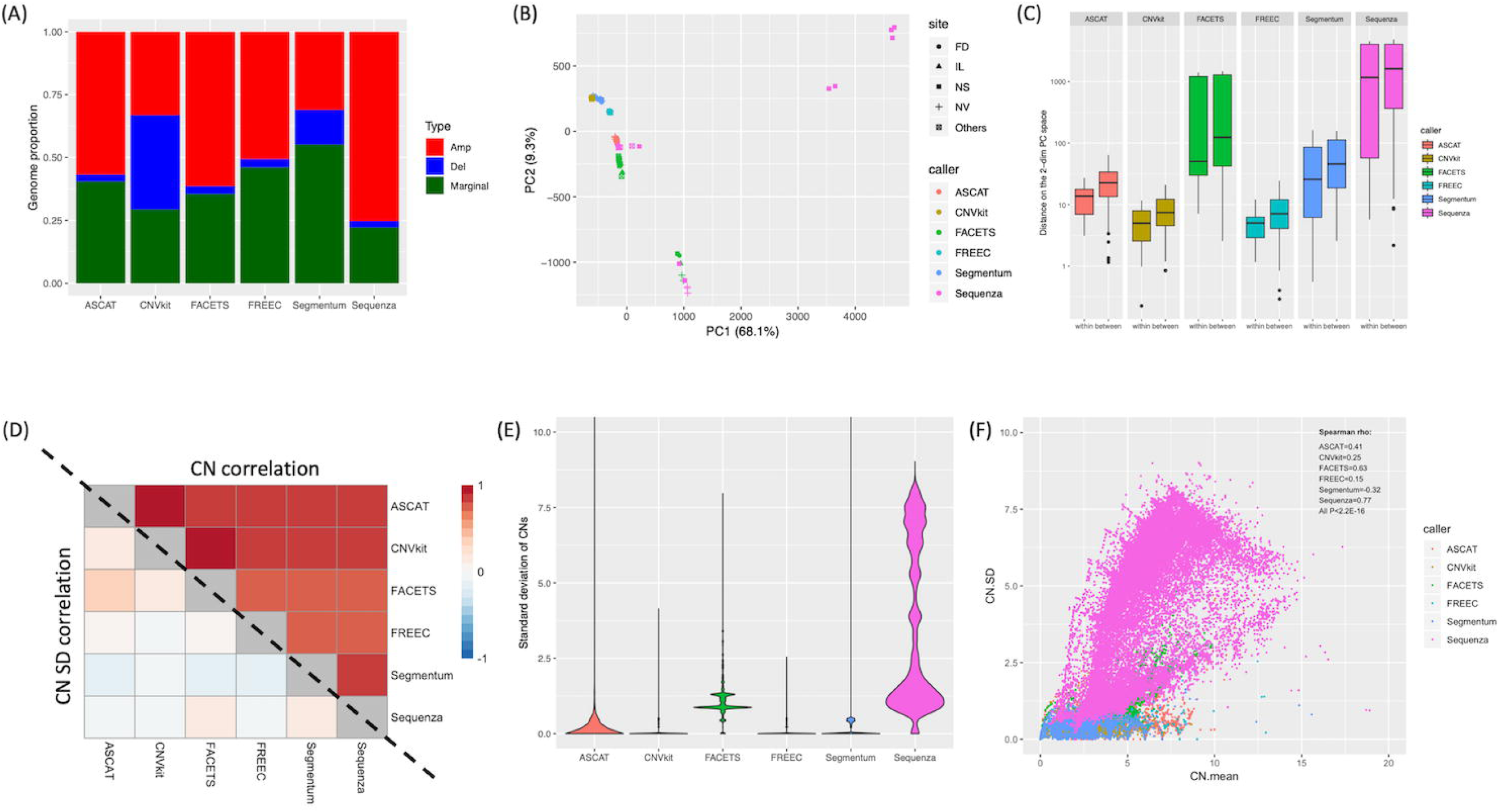
Variation of CNVs across the 21 reference (REF) samples. All analyses were based on the binned genome (see Methods for how to compute copy number (CN) for each bin using the raw CNV calls). (A) Proportion of genome called as amplification (red), deletion (blue), and marginal (dark green) based on the median CNs estimated by each caller across the REF samples. (B) Principal component analysis for 126 (21 x 6) sets of CNV calls. (C) Distribution of within-site and between-site pairwise distances in (B) for each caller. Between-site distances were significantly higher than within-site distances (Wilcoxon rank sum test; P<0.05) for all callers. (D) Spearman correlation of median CNs (the upper-triangular matrix) and standard deviation (SD) of CNs (the lower-triangular matrix) across REF samples between callers. (E) Distribution of SDs of CNs across the REF samples. Width of the violin plot is proportional to the number of bins. (F) The scatter plot for mean of CN (x-axis) versus SD of CN (y-axis). Spearman correlation is shown at top-right corner.

To evaluate consistency of CNV calls, we computed standard deviation (SD) of CNs across 21 REF replicates for each bin for each caller. CNV calls generated by CNVkit, FREEC, and Segmentum were very consistent across replicates (SD of 0 for most bins) whereas Sequenza showed the highest CN variation (Figure 2E). The CN SD was positively correlated with CN mean in all callers except Segmentum (Figure 2F), suggesting the estimated CNs were not reliable if they were too high. In addition, we wondered if callers consistently produced variable CNs in particular genomic regions (bins). We computed (Spearman) correlation of CN SDs across bins between callers but no strong correlation was seen overall (Figure 2D). The mild positive correlation was likely driven by those regions with high CNs where CNs tended to be varied among replicates as shown in Figure 2F. Given the low correlation of CN SDs between callers, those bins with variable CNs were mainly caller-specific but not a systematic issue associated with NGS-based CNV calling.

### CNV calling drastically impacted by low tumor purity

Next, we investigated effects of 6 major experimental and technical confounding factors: medium tumor purity (50% and 75%), low tumor purity (<50%), FFPE-processed (versus fresh) samples, low input DNA amount (1ng and 10ng), WES (versus WGS), and low sequencing coverage (10X and 30X with 100% tumor purity). To avoid the analysis being confounded by the choice of callers (which have been shown above to have huge impact on CNV calls), we compared the 70 confounded samples (affected by different confounding factors) to the 21 REF samples within each caller. To quantify variation and discrepancies of CNV calls between confounded samples and REF samples, we performed PCA on all 91 samples (Supplementary Figure 1) and computed the factor-to-REF distance (Euclidean distances between median CNs of confounded samples and those of REF samples) and the within-condition distance (mean pairwise distance within each condition including REF). A large factor-to-REF distance indicates pronounced changes in CNV calls introduced by the confounding factor (compared to REF) whereas a large within-condition distance indicates the large variation of CNV calls across replicates within a particular condition. We observed that low tumor purity greatly affected CNV calling because it consistently produced large factor-to-REF distances among 5 confounding factors (Figure 3A) and low purity samples were visually away from the REF samples in PC plots for all callers (Supplementary Figure 1). Medium purity, WES and FFPE also showed pronounced impacts on CNV calling whereas impacts of, low coverage and low input DNA amount were relatively mild (Figure 3A). ASCAT, FACETS, and Sequenza displayed higher variation across replicates in almost all conditions (Supplementary Figure 2A). Although variation in CNVkit, FREEC, and Segmentum was mild (Supplementary Figure 2A), their variation was clearly increased by confounding factors such as medium purity, WES, and FFPE as evidenced by the ratio of within-condition distances between the confounding factor and REF greater than 1 (Supplementary Figure 2B).

**Figure 3.**
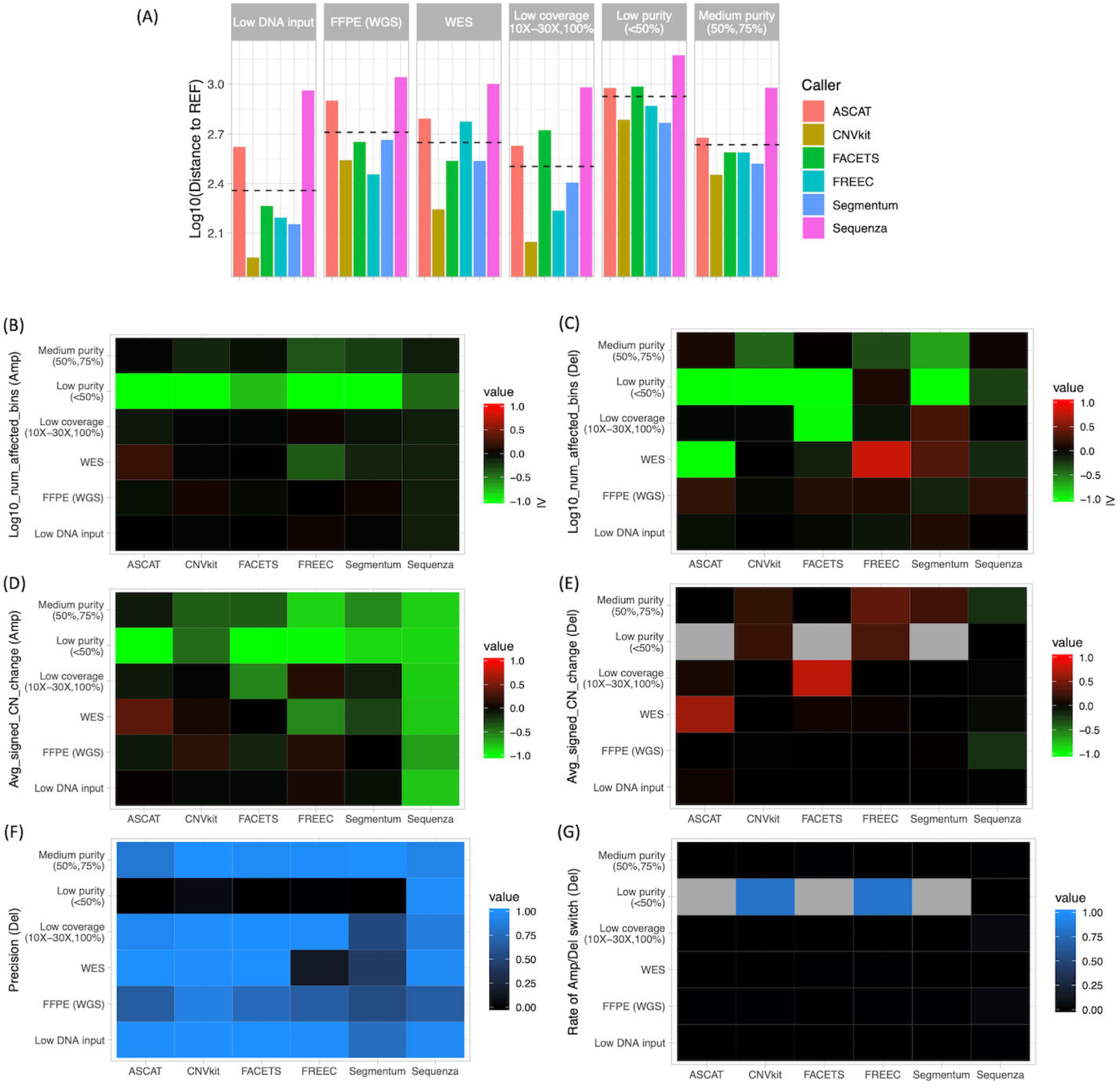
Effects of the confounding factors on CNV calls. Six confounding factors (y-axis) were analyzed for each of the 6 callers (x-axis) in each figure: medium purity (purity level=50% and 75%), low purity (purity level < 50%), low coverage (10X-30X with 100% purity), WES (versus WGS), FFPE-processed samples (versus fresh samples), and low input DNA amount. Analyses were done within each caller by comparing median CNs between two groups: the samples confounded by a particular factor and the reference (REF) samples. (A) Euclidean distance between two groups (in Log base 10). (B and C) Ratio of the number of (B) amplification and (C) deletion bins between two groups (in Log base 10). (D and E) Average sign change (+1 if CN increased, -1 if decreased, and 0 if not changed) across bins for (D) amplification and (E) deletion calls. (F) Precision of deletion calls when taking REF as the truth set. (G) Fraction of bins called as deletion in the confounded samples but called as amplification in the REF samples. Note that all analyses were based on the binned genome (see Methods for how to compute copy number for each bin using the raw CNV calls).

### Low tumor purity missed many CNV calls, greatly changed CNs, and called deletions incorrectly while effects of other confounding factors were caller-specific

To investigate how CNV calls were affected, the median CNs (across replicates) were used for each confounding factor and REF in the following analyses. First, we computed the ratio of numbers of amplified (and deleted) bins between each confounding factor and REF. A large ratio (>1) indicates more amplification (or deletion) bins compared to REF whereas a small ratio (<1) indicates fewer bins meaning that some bins called in REF were missed when samples were confounded by particular factors (Figures 3B and 3C). For both amplification and deletion, low tumor purity consistently produced small ratios in all callers (except FREEC for deletion) indicative of many amplified and deleted bins being missed. The same trend was seen for medium purity in CNVkit, FREEC, and Segmentum but less intensive. WES produced more amplified and fewer deleted bins when using ASCAT but fewer amplified and more deleted bins when using FREEC and Segmentum. On the other hand, FFPE tended to generate more deleted bins.

Next, we computed the average sign change (+1 if CN was larger in the confounding factor versus the REF, 0 if CN did not change, and -1 if CN was smaller) across those bins called as amplification (or deletion) in both the particular confounding factor and the REF (Figures 3D and 3E). The positive sign change indicates overall increase of estimated CNs for bins called as amplification (or deletion) while the negative sign change indicates decrease of estimated CNs. The sign change was close to -1 for amplification across all callers (except CNVkit which was - 0.45) for low tumor purity, showing that even for those bins remained called as amplification, their CNs were estimated lower when tumor purity was low. The effect of low tumor purity on deletion bins was opposite – increasing CNs for deleted bins. The same trend was also seen for medium purity in FREEC and Segmentum but less intensive. In addition, WES increased CNs for both amplified and deleted bins using ASCAT but decreased CNs using FREEC for amplified bins. When using FACETS, low coverage decreased CNs for amplified bins but increased CNs for deleted bins. Sequenza seemed sensitive to all confounding factors as evidenced by its consistent decrease of CNs for amplified bins for all factors.

Lastly, we wondered if CNV calls were called correctly in confounded samples. We computed the precision (i.e. the proportion of amplified (or deleted) bins called in a confounding factor also being called as amplification (or deletion) in the REF) and the fraction of switching calls (i.e. the proportion of amplified (or deleted) bins in a confounding factor being called as deletion (or amplification) in the REF). Precision was high for amplification calls overall and no switching call was seen even when tumor purity was low except WES displayed a little low precision when using ASCAT (Supplementary Figures 2C and 2D). However, deletion calls were problematic under low tumor purity where deleted bins were either not called or called as amplification in the REF (Figures 3F and 3G). Sequenza seemed the only one which was not affected much in terms of precision of calling deletion in low purity samples. The precision of deletion calls was generally decreased when samples were FFPE-processed and when Segmentum was used. In addition, the precision of deletion calls dropped drastically using FREEC when WES was performed.

In summary, low tumor purity greatly affected CNV calling by missing most calls, lowering (increasing) CNs for amplification (deletion) calls, and producing incorrect deletion calls. Medium purity mainly affected CNVkit, FREEC, and Segmentum in the similar trend as low purity did but with less intensity. FFPE-processed samples increased variability and lowered precision of CNV calls. Compared to WGS, WES affected CNV calls by increasing variability, missing or producing incorrect calls depending on callers, and lowering the precision of deletion calls. Effects of low input DNA amount and low coverage were relatively minor. Effects of the 5 confounding factors on all 6 callers were summarized in Table 1. Supplementary Table 2 contains all data used to plot Figure 3 and Supplementary Figure 2. Supplementary Figure 3 shows the overview of amplification and deletion calls by all callers in all conditions (including confounding factors and REF).

**Table 1.** Summary of impacts of confounding factors on CNV calling.

### Accuracy of purity estimation varied at low purity

Among the 6 callers, ASCAT, FACETS, and Sequenza explicitly modeled sample tumor purity in CNV calling. Because low tumor purity was identified as the major confounding factor in our analysis, we wondered if the accuracy of tumor purity estimates was correlated with CNV calling quality. For each low purity sample (tumor purity = 5%, 10%, and 20% with coverage >= 50X), we measured the calling quality as the Euclidean distance between its CNs and the median CNs across the REF samples (the lower, the better) and computed the absolute difference between the true purity and its purity estimate as the purity estimation accuracy (the lower, the better). We found the calling quality was indeed positively correlated with the purity estimation accuracy for ASCAT (Spearman rho=0.6, P=0.02) and Sequenza (rho=0.5, P=0.06), suggesting that accurate purity estimation for low purity samples could achieve better calling quality. FACETS was not included in the calculation because many of its purity estimates were not available.

Next, we asked whether tumor purity could be estimated accurately by computational methods. Five methods were included in this analysis: 4 CNV callers, ASCAT, FACETS, PureCN [26], and Sequenza, in which tumor purity is explicitly modeled in the calculation, and THetA2 [27], a method developed for tumor purity estimation. Purity was estimated for the 42 samples with varying purity levels and sequencing coverages (Figure 1A). We observed that the absolute error (absolute difference between the true purity and the estimate) was clearly higher for purity of 5% and displayed less consistency (larger variance and outliers) at low purities (5%, 10%, and 20%) (Figure 4A). Sequencing coverage, however, did not affect purity estimation much except a little higher variation of performance was introduced at coverage of 10X. The RMSE values for Sequenza, PureCN, ThetA2, FACETS, and ASCAT were 0.095, 0.108, 0.181, 0.233, and 0.348 respectively, suggesting that Sequenza and PureCN outperformed other tools in our analysis. Specifically, ASCAT dramatically overestimated purities of 5%, 10% and 20% except 200X at 20%. ThetA2 tended to underestimate at high purity levels (75% and 100%) (Figure 4B). FACETS often failed to estimate purity at 5%. Raw purity estimates were reported in Supplementary Table 3.

**Figure 4.**
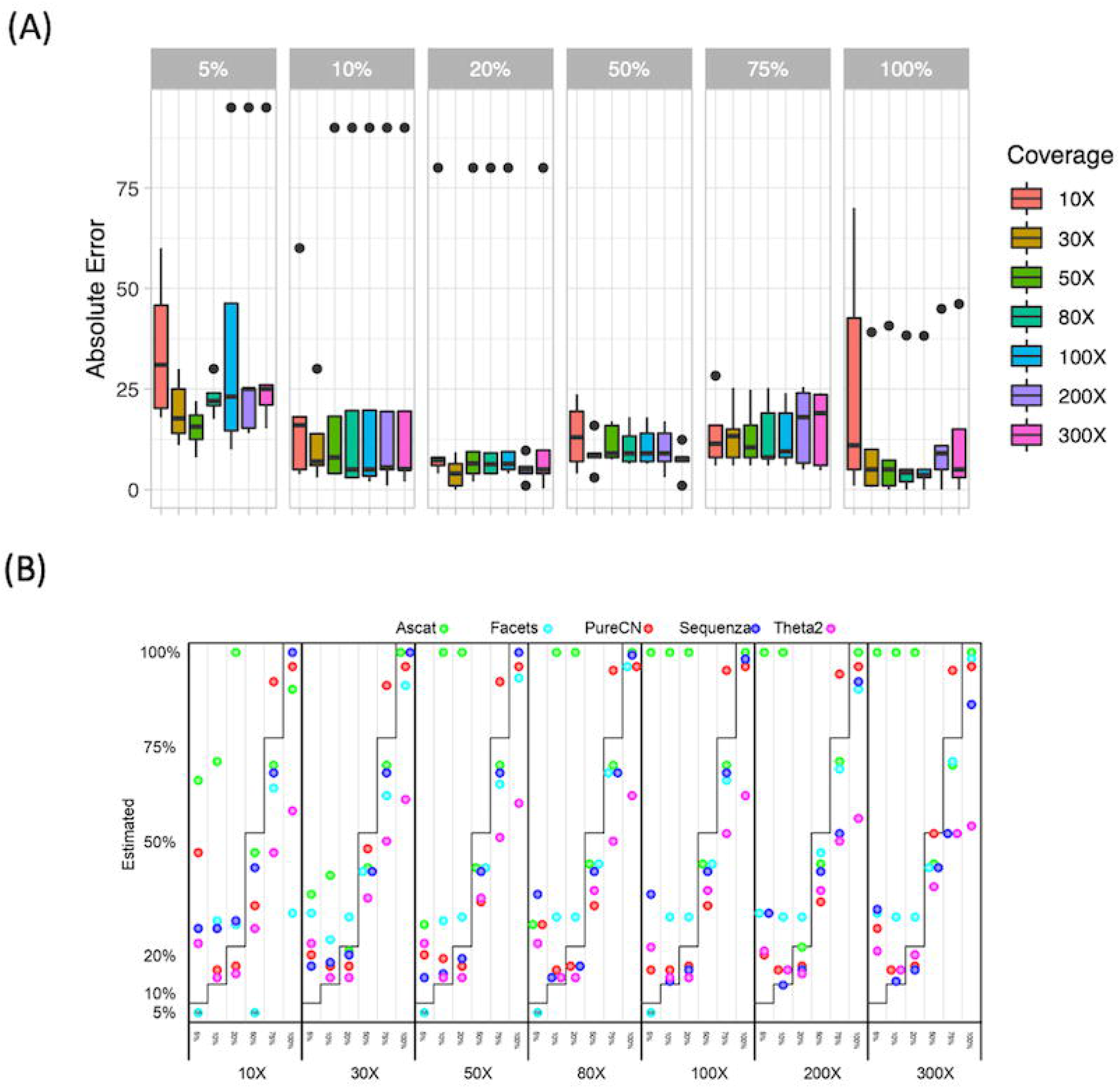
Purity estimation of five computation tools at different purity levels and sequencing coverages. (A) Distribution (across the five tools) of absolute differences between the true purity and purity estimates. (B) Purity estimates reported by the five computation tools.

### CNVkit and Segmentum performed better when benchmarked using the array-based calls but worse using majority calls and a set of high confidence calls

To evaluate performance of the 6 callers, we focused on the REF samples. We first used an independent set of CNV calls generated by a non-NGS-based technology as the truth set. We performed Affymetrix CytoScan Array, which incorporates 2.68 million genomic markers to detect genome-wide chromosomal abnormalities [28, 29] and categorized each bin into amplification, deletion, or marginal as we did for NGS-based callers. Overall, array calls were more similar to CNVkit and Segmentum (Figure 5A). We benchmarked our CNV callers by calculating sensitivity, specificity, precision, and F1 score for amplification, deletion, and marginal (Figure 5B and Supplementary Figure 4). Not surprisingly, overall, Segmentum and CNVkit outperformed the other 4 callers with higher F1 scores for both amplification and deletion. We also noted that many deletions called by CNVkit and Segmentum were called marginal by the other 4 callers while some marginal calls called by CNVkit and Segmentum were called amplifications by the others (Figure 5A). This was consistent with our previous observation that despite much disagreement in the proportion of amplification and deletion between callers, their CNs were highly correlated (the upper triangular matrix in Figure 2D).

**Figure 5.**
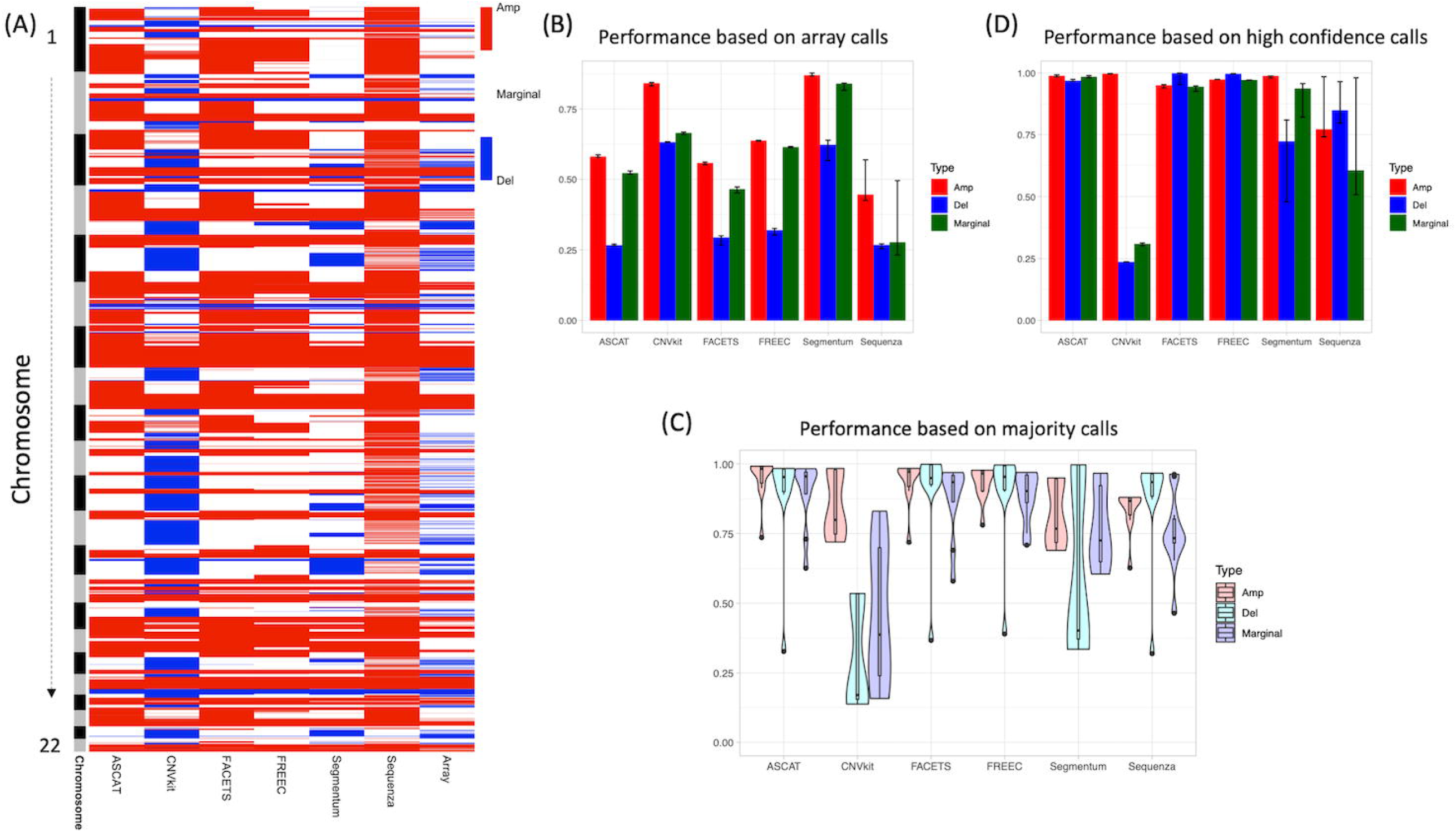
Performance evaluation for callers. (A) Amplification (red), deletion (blue), and marginal (white) calls across the entire genome called by the NGS-based callers (names labeled at bottom) and Affymetrix CytoScan Array. The row side bar on the left indicates chromosomes ordered from 1 (top) to 22 (bottom). (B) F1 score (y-axis) for the 6 callers (x-axis) computed for amplification (red), deletion (blue), and marginal (dark green) calls using array calls as the truth set. The bar shows the median F1 across the 21 reference (REF) samples and the error bar indicates 25% and 75% percent quantile. (C) Distribution of F1 scores (y-axis) for the 6 callers (x- axis) computed for amplification (light red), deletion (light blue), and marginal (light purple) calls using majority calls as the truth set. The majority calls were the CNVs called in at least 2 callers out of the 3 (based on their median copy numbers across the 21 REF samples). Evaluation was done for each of 10 (5-choose-2) combinations for all 21 REF samples, totaling 210 F1 scores. The box plot indicates 10%, 25%, 50%, 75%, and 90% quantiles. Note that all analyses were based on the binned genome (see Methods for how to compute copy number for each bin using the raw CNV calls). (D) F1 score (y-axis) for the 6 callers (x-axis) computed for amplification (red), deletion (blue), and marginal (dark green) calls using a set of high confidence calls as the truth set. The bar shows the median F1 across the 21 reference (REF) samples and the error bar indicates 25% and 75% percent quantile.

Majority rule is often used to generate the high confidence call for evaluating computational tools when the real ground truth cannot be obtained [30, 31]. In practice, 3 callers (including the one being evaluated) would be applied to the same data and those calls which are agreed in at least 2 callers would be taken as the ground truth. Evaluation was done using median CNs across the 21 REF samples. For each caller, evaluation was performed 10 times to cover all combinations (choosing 2 out of the other 5 callers). For each combination, the majority calls were those calls agreed in at least 2 out of 3 callers (the 2 chosen ones plus the one being evaluated). As expected, ASCAT, FACETS, FREEC, and Sequenza performed better overall in terms of F1 scores (Figure 5C) because they formed the majority among the 6 callers and more agreed to each other compared to CNVkit and Segmentum (Figure 5A). However, F1 scores varied depending on which 2 callers were chosen to generate majority calls. When CNVkit and Segmentum were included in the evaluation, more deletions were generated in the majority calls. Variation was also seen for precision, sensitivity, and specificity (Supplementary Figure 5). Performance varied even larger if sample variation was taken into account (i.e. using CNs of a randomly picked REF sample for each caller instead of taking median CNs across the 21 REF samples) (Supplementary Figure 6).

The third approach was to evaluate callers using a set of high confidence calls. Rather than using all bins across the whole genome, we restricted the evaluation within 134,786 out of 287,509 (46.88 %) bins where CNV calls (i.e. amplification, deletion, or marginal) agreed across 4 out of 6 callers and array calls, including 63,379 marginal, 63,451 amplification and 7,956 deletion bins (Supplementary Table 4). In terms of F1 score, ASCAT, FACETS, and FREEC performed better compared to others whereas CNVkit performed poorly because it called many marginal bins as deletion within this set of high confidence calls (Figure 5D; precision, sensitivity, and specificity shown in Supplementary Figure 7).

### Precision of CNV calling can be improved by taking consensus calls or calling CNVs using stringent cutoffs

In practice, because obtaining the true set of CNV calls across the genome is almost impossible in cancer studies, rather than reporting many CNV calls with unknown (possibly low) accuracy, we often prefer reporting fewer calls with high confidence (high precision). Thus, we wondered whether (and how) we can possibly improve the precision of CNV calls in NGS-based CNV analyses. We used the array-based CNV calls as the truth set in this analysis. First, we investigated if CNVs called by more than one caller (n>1; i.e. n=2, 3, 4, 5, or all 6 callers) would achieve higher precision. For each n, we computed the average precision over all combinations. For example, the precision for n=2 was averaged over 15 (6 choose 2) combinations. Then, we checked if (and how much) the precision of consensus calls for n callers was improved compared to the best precision among those for less-than-n (including n=1) callers (Figure 6A). We observed that precision went higher when taking consensus calls across 2 callers regardless of the influences of confounding factors. The increase was minimal when n=3 and saturated for n>=4. For low purity samples, consensus calls might decrease precision or result in no call. However, taking consensus calls across more callers would lower sensitivity and, thus, might lower the F1 score as precision increased (Supplementary Figure 8A).

**Figure 6.**
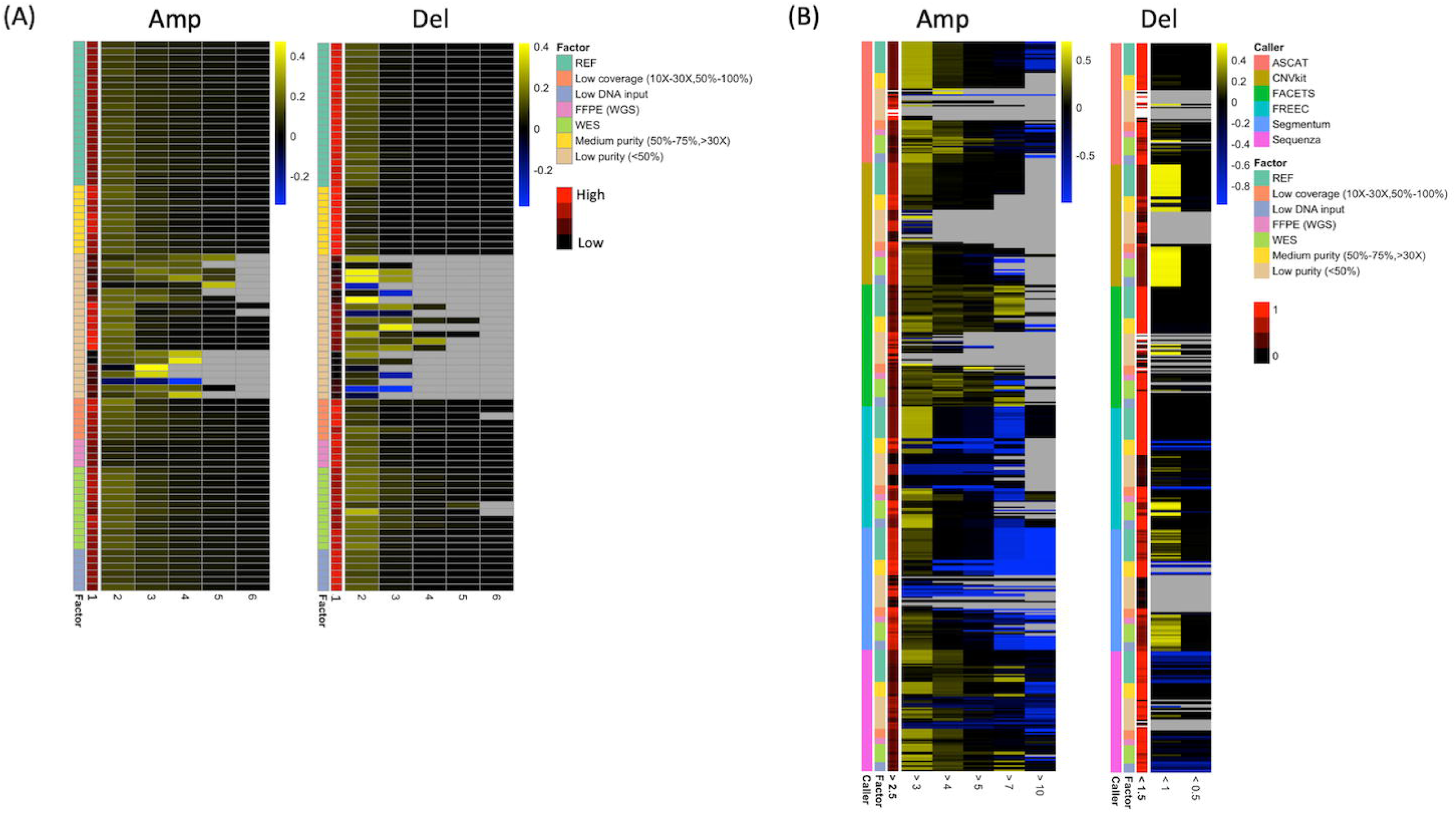
Precision of CNV calls can be improved by taking consensus calls or applying stringent calling cutoffs. (A) Heatmaps for amplification (left) and deletion (right) calls. Two side bars on the left indicate 7 sample conditions (in different colors) and average precision across the 6 tested CNV callers (in red scale). Heatmaps show precision increase when taking consensus calls across n=2 to 6 callers (from left to right). Precision increase: the difference between the precision at n=k and the best precision among n=1 (indicated in the row side bar in red scale) to k-1. (B) Heatmaps for amplification (left) and deletion (right) calls. Three side bars (from left to right) indicate 6 callers, 7 sample conditions and precision (in red scale) for the original cutoff used in this study (>2.5 for amplification and <1.5 for deletion). Heatmaps show precision increase by applying more stringent cutoffs (labeled at bottom) to call CNVs. Precision increase: the difference between the precision for a specific cutoff and the best precision among less stringent cutoffs. Note that all analyses were based on the binned genome (see Methods for how to compute copy number for each bin using the raw CNV calls).

Next, we tested if applying stringent cutoffs for calling amplification and deletion can improve precision. In this study, we called amplification and deletion by CN >2.5 and CN <1.5 respectively. Here, we tried to vary cutoffs (amplification: >3, >4, >5, >7, and >10; deletion: <1 and <0.5) and computed precision for each cutoff. We found applying CN>3 for amplification and CN<1 for deletion increased precision (Figure 6B). More stringent cutoffs did not increase precision much further (e.g. CN>4 and CN>5 for amplification and CN<0.5 for deletion) and too stringent cutoffs could impair precision instead (e.g. CN>7 and CN>10 for amplification). As expected, the F1 score decreased as the cutoff went more stringent except applying CN>3 for amplification where the F1 score was higher than applying CN>2.5, suggesting that a slightly stringent cutoff for amplification (i.e. CN>3) could be a good choice of obtaining high precision without scarifying much sensitivity (Supplementary Figure 8B).

We summarized results of our analyses in Table 2 for all 6 callers including variation across replicates, proportion of amplification and deletion calls, performance, effects of confounding factors, and purity estimation.

**Table 2.** Summary for all 6 NGS-based CNV callers.

## Discussion

To our knowledge, this study presents the most comprehensive assessment of NGS-based genome-wide somatic CNV detection using real sequencing data. We systematically investigated how reproducibility of CNV detection is affected by experimental and technical confounding factors including impure tumor samples, FFPE-processed (versus fresh) samples, low input DNA amount, WES (versus WGS), and low sequencing coverage as well as sources of variability such as sequencing sites and CNV callers. When no confounding factor was involved, the variability of CNV calls was mainly due to callers. ASCAT, FACETS, and Sequenza displayed higher CNV calling variation across replicates. Within each caller, high variation often correlated with highly amplified regions. Low tumor purity (<= 20%) was the major confounding factor, making purity estimation unreliable and drastically affecting CNV calling in 3 aspects: (1) many CNV calls were missed; (2) CNs of affected regions were changed toward marginal (lower CN for amplification and higher CN for deletion); and (3) precision of deletion calls was impaired. WES, FFPE-processed samples, and samples with 50% - 70% tumor purity increased variation of CNV calls across replicates but affected the calls in a caller-specific manner. Effects of low input DNA amount and low coverage were relatively minor. CNVkit and Segmentum outperformed others when benchmarking using CNV calls produced by the non-NGS-based cytogenetic array whereas ASCAT, FACETS, and FREEC performed better when the majority calls among the 6 NGS-based callers and a set of high confidence calls were used for evaluation. Finally, we found taking consensus calls across multiple callers and applying stringent cutoffs for calling amplification and deletion could improve precision of CNV calls.

The comprehensive analysis conveys important messages for NGS-based CNV analysis in cancer. First, we recommend that rather than using the reported CNs for downstream analysis, sorting CNV-affected regions into categories (such as amplification, deletion, and marginal) can reduce variation caused by unknown factors because the reported CNs could be unreliable especially for those regions with high CNs. Second, preparing fresh samples followed by WGS is preferred but ensuring high tumor purity is critical to have reliable CNV calls. If tumor purity is less than 50%, we recommend interpreting deletions cautiously because they could be either marginal or amplification. Third, because variability of CNV calls is dominated by CNV callers, to increase confidence, we recommend taking consensus calls across multiple callers (e.g. n=2 or 3) and using slightly more stringent cutoffs to call amplifications (e.g. CN>3) and deletions (e.g. CN<1.5 or CN<1). However, taking consensus calls across too many callers (e.g. n>=4) or using too stringent cutoffs (e.g. CN>=4 for amplification and CN<0.5 for deletion) may greatly increase false negatives or even decrease precision (e.g. CN>7 for amplification in our analysis). Finally, given the high variability of CNV calls, in addition to the approaches mentioned above, we recommend validating important CNV findings in replicates or independent datasets or gaining support from other evidences such as FISH and differential expression of affected genes.

Our study highlights the challenge of benchmarking methods for analyzing heterogeneous cancer data when the real ground truth is not available. In the past, developers of CNV callers often demonstrated superior performance of their methods by using an independent set of calls generated by non-NGS-based technologies (such as FISH or arrays) or using majority calls reported by other existing callers as the ground truth. However, we showed that in this study, array calls favored CNVkit and Segmentum whereas majority calls favored the other 4 callers, indicating that evaluating method performance using either approach could be biased. Large variation in resulting performances using majority calls can be introduced by the choice of tools for creating majority calls especially when CNV calls reported by available tools fall into very inconsistent categories (e.g. ASCAT, FACETS, FREEC, and Sequenza in one category versus CNVkit and Segmentum in another category in this study; see Figure 2A and Figure 5A). When restricting performance evaluation within a set of high confidence calls, CNVkit and Sequenza showed producing more false positives for deletion and amplification respectively while ASCAT, FACETS, and FREEC were more reliable. Lacking the real ground truth will likely remain an issue in the near future for cancer data analysis. To be unbiased, we recommend method developers to evaluate their methods in multiple ways and present all the results to give potential users a comprehensive view of the newly developed methods. To be robust, we recommend data analysts to validate their findings. For example, one can validate important CNV findings using FISH or checking if consistent results are seen in other data types (e.g. no expression in RNA-seq data is seen when the gene is predicted as homozygous deletion).

Although method evaluation studies often report the best method as their final conclusion, picking such caller for somatic CNV calling seems currently impractical in this study. In addition to the limitation of lacking ground truth for performance evaluation, none of the callers performed consistently better in all aspects (including situations where confounding factors existed). For example, Sequenza was the best in purity estimation (Figure 4B) and indeed displayed less sensitive to low purity compared to other tools (Figures 3B, 3C, and 3E). However, the variation of its CNV calls was the highest (Supplementary Figure 2A), indicative of its sensitivity to stochastic noise. CNVkit and FREEC had the lowest variation (Supplementary Figure 2A) but CNVkit became an outlier calling the most genome deletion (Figure 5A) while FREEC was more affected by WES (Figures 3B, 3C, 3D, and 3F). ASCAT and FACETS showed a medium level of variation and were affected by WES and low coverage respectively. Segmentum also displayed low variation in CNV calls but it seemed sensitive to all confounding factors as sensitivity of deletion calls dropped in impure samples and precision dropped when other factors were introduced. Therefore, we believe it would be more practical to fairly present strength and weakness of each caller (a comprehensive view of caller characteristics summarized in Table 2) and to provide guidelines to improve CNV calling precision (e.g. using consensus calls and applying slightly stringent cutoffs) rather than picking the best caller. Researchers can pick the caller suitable for their experimental conditions and increase confidence of their findings using computational strategies.

ASCAT, FACETS, and Sequenza explicitly model tumor purity in CNV calling whereas CNVkit, FREEC, and Segmentum do not. Although none of them performed well for low purity (<50%) samples, ASCAT, FACETS, and Sequenza did perform better for samples with 50% and 75% purity as evidenced by no variation increase relative to REF (Supplementary Figure 2B), less calls being missed (Figures 3B and 3C), and milder CN changes (Figure 3E). Purity estimation analysis also showed lower estimation error and more consistent estimates at purity >= 50% (Figure 4A). Because most tumors were reported with purity > 50% in TCGA across all cancer types [32], low purity (<50%) is likely prevented by attentive sample collection. This strengthens the importance of modeling tumor purity in somatic CNV calling since the majority of tumor samples would be neither perfectly pure nor with very low purity (<50%). However, we suspect that modeling tumor purity may make callers sensitive to stochastic noise due to model complexity as we observed that ASCAT, FACETS, and Sequenza coincided with higher variation across replicates (Supplementary Figure 2A) and higher CN-SD correlation (Figure 2F). In light of these observations, in the future, it would be a pressing need to develop computational methods that can model tumor purity and be robust to data noise for somatic CNV calling.

In conclusion, we present a comprehensive study for somatic CNV calling using real sequencing data. We have analyzed the source of variation, investigated robustness and effects of confounding factors of CNV calling for cutting-edge callers, discussed the bottleneck of method evaluation in somatic CNV calling, and explored practical guidelines for improving confidence of CNV calls. A list of observations and recommendations was summarized in Box 1. We hope our analyses would not only help researchers to optimize their experimental design and CNV analysis but also guide method developers to improve the CNV calling algorithm in the future.

## Methods

### Experimental design and initial analysis

We used the SEQC-II reference matched samples, a human triple-negative breast cancer cell line (HCC1395) 291 and a matched B lymphocyte derived normal cell line (HCC1395BL) in our analysis. Detailed sample information as well as DNA extraction, FFPE processing and DNA extraction, DNA fragmentation and library preparation, DNA sequencing, read processing and quality control, read alignment, and GATK indel realignment and quality score recalibration can be found in the SEQC-II reference samples manuscripts [24, 25].

### Assign copy number (CN) for each bin

The entire genome is binned into 287,509 bins (bin size = 10k) with a normal CN = 2. If a bin_i_ is covered by any CNV calls > 1kb, CN of bin_i_ is the weighted (length of overlapping regions) mean of CNs called. A bin is considered as amplification or deletion if the CN is > 2.5 or < 1.5 respectively. In addition to using the fixed-length bins as the units for CNV calls, we also computed CNs using genes as the units. The gene-based approach was computed the same way as the bin-based approach. The proportion of amplification and deletion based on the gene-based approach was pretty much the same as those based on the bin-based approach (Supplementary Figure 9), suggesting our choice of bin size did not affect analysis much.

### Tumor purity estimation

We used five purity inference tools, ASCAT (AscatNgs 4.2.1), FACETS (0.6.0), PureCN (1.12.2), THetA (0.7) and Sequenza (2.1.2), which are all available on the NIH Biowulf HPC cluster. All tools are run using default parameters or parameters recommended by the user’s manual. More specifically, for AscatNgs, we added flag “-protocol WGS”. For FACETS, we used “snp-pileup” and set “cval=50 for procSample”. For Sequenza, we used pileups as input files and ran seqz_binning with “-w 100”. For PureCN, we included flags “--funsegmentation none --force -- postoptimize --seed 123” and for THetA, we used the updated algorithm THetA2 with flags “-n 2 -k 4”. Segment files from CNVkit (0.9.1 --method wgs) were used as input files for both PureCN and THetA. For tool performance ranking, we used root mean square error (RMSE) to compare the estimates to the truth purity for each condition and then calculated the RSME mean for final ranking.

## Data and Code Availability

The raw and aligned data could be obtained via Fang et al [24]. The consensus sets of CNVs as well as CNV callers’ scripts and setting are available on ftp://ftp-trace.ncbi.nlm.nih.gov/ReferenceSamples/seqc/.

## Acknowledgements

This study utilized the high-performance computational capabilities of the Biowulf Linux cluster at the National Institutes of Health, Bethesda, MD.

## Disclaimer

This is a research study, not intended to guide clinical applications. The views presented in this article do not necessarily reflect current or future opinion or policy of the National Institutes of Health and or US Food and Drug Administration. Any mention of commercial products is for clarification and not intended as endorsement.

## Author contributions

DM, MP, and WX conceived the study; CN, FS, and MP set up and ran CNV callers; CN, CY, QC, and DM performed tumor purity analysis; YC designed and implemented analyses of CNV calls; MP and YC wrote the manuscript; CW and WX coordinated sample preparation and collection; DM, MP, and WX supervised the project; All authors reviewed and edited the manuscript.

## Supplementary Figures

**Supplementary Figure 1. Within-caller principal component analysis for the reference samples and samples confounded by different factors.** Principal component (PC) plots (the 1^st^ PC on x-axis versus the 2^nd^ PC on y-axis) for each caller (caller name labeled at top-right corner). Note that all analyses were based on the binned genome (see Methods for how to compute copy number for each bin using the raw CNV calls).

**Supplementary Figure 2. Effects of the confounding factors on CNV calls.** Six confounding factors (y-axis) were analyzed for each of the 6 callers (x-axis) in each figure: medium purity (purity level=50% and 75%), low purity (purity level < 50%), low coverage (10X-30X with 100% purity), WES (versus WGS), FFPE-processed samples (versus fresh samples), and low input DNA amount. (A) The heatmap shows the mean of sample pairwise distances (on the PC spaces shown in Supplementary Figure 1) within each condition (including the reference (REF)). (B) The heatmap shows the ratio of mean pairwise distances between each confounding factor and the REF in (A) (i.e. values in each row divided by values in the 1^st^ raw in (A)). A ratio > 1 (red) indicates increase of variation in a specific factor compared to the REF whereas a ratio < 1 (green) indicates decrease of variation. (C) Precision of amplification calls. (D) Fraction of CNVs called as amplification in confounded samples but called as deletion in the REF samples. Note that all analyses were based on the binned genome (see Methods for how to compute copy number for each bin using the raw CNV calls).

**Supplementary Figure 3. Overview of CNVs called by each of the 6 callers for all samples (confounded + reference).** The heatmap shows amplification (red), deletion (blue), and marginal (white) calls across the entire genome called by the NGS-based callers for all samples. Each row is a fixed-length genomic region and each column presents the CNVs called by a specific caller for a specific sample. The row side bar on the left indicates chromosomes ordered from 1 (top) to 22 (bottom). The column side bars indicate the confounding factors and the callers.

**Supplementary Figure 4. Performance evaluation for the 6 NGS-based callers using CNVs called by Affymetrix CytoScan Array.** (A) Sensitivity (y-axis), (B) Specificity (y-axis), and (C) Precision (y-axis) for the 6 callers (x-axis) computed for amplification (red) and deletion (blue) calls using array calls as the truth set. The bar shows the median across the 21 reference (REF) samples and the error bar indicates 25% and 75% percent quantiles.

**Supplementary Figure 5. Performance evaluation for the 6 NGS-based callers using majority calls.** Distributions of (A) Sensitivity (y-axis), (B) specificity (y-axis), and (C) precision (y-axis) for the 6 callers (x-axis) computed for amplification (light red), deletion (light blue), marginal (light purple) calls using majority calls as the truth set. The majority calls were the CNVs called in at least 2 callers out of the 3 (using their median copy numbers across the 21 REF samples). Evaluation was done for each of 10 combinations (5 choose 2) for all 21 REF samples, totaling 210 scores. The box plot indicates 10%, 25%, 50%, 75%, and 90% quantiles. Note that all analyses were based on the binned genome (see Methods for how to compute copy number for each bin using the raw CNV calls).

**Supplementary Figure 6. Performance evaluation for the 6 NGS-based callers using a set of high confidence CNV calls.** (A) Sensitivity (y-axis), (B) Specificity (y-axis), and (C) Precision (y-axis) for the 6 callers (x-axis) computed for amplification (red), deletion (blue), marginal (dark green) calls using array calls as the truth set. The bar shows the median across the 21 reference (REF) samples and the error bar indicates 25% and 75% percent quantiles.

**Supplementary Figure 7. Variation of performance evaluated by the majority calls increased when sample variation was introduced.** (A) F1 score (y-axis), (B) sensitivity (y-axis), (C) specificity (y-axis), and (D) precision (y-axis) for the 6 callers (x-axis) computed for amplification (light red) and deletion (light blue) calls using majority calls as the truth set. They were computed the same way as Figure 5C and Supplementary Figures 5A, 5B, and 5C respectively, except the majority calls were generated across randomly picked reference (REF) samples rather than the median copy numbers (among the 21 REF samples) for the 3 chosen callers.

**Supplementary Figure 8. F1 score of CNV calls may or may not be improved by taking consensus calls or applying stringent calling cutoffs.** (A) Heatmaps for amplification (left) and deletion (right) calls. Two side bars on the left indicate 7 sample conditions and average precision across the 6 tested CNV callers (in red scale). Heatmaps show F1 increase by taking consensus calls across n=2 to 6 callers (from left to right). F1 increase: the difference between the F1 at n=k and the best F1 among n=1 (indicated in the row side bar in red scale) to k-1. (B) Heatmaps for amplification (left) and deletion (right) calls. Three side bars (from left to right) indicate 6 callers, 7 sample conditions and F1 (in red scale) for the original cutoff used in this study (>2.5 for amplification and <1.5 for deletion). Heatmaps show F1 increase by applying different cutoffs (labeled at bottom) to call CNVs. F1 increase: the difference between the F1 for a specific cutoff and the best F1 obtained from less stringent cutoffs. Note that all analyses were based on the binned genome (see Methods for how to compute copy number for each bin using the raw CNV calls).

**Supplementary Figure 9. Proportion of genome called as amplification, deletion, and marginal computed for each caller using the gene-based approach (see Methods).**

## Supplementary Tables

**Supplementary Table 1. Copy numbers of 287,509 bins (bin size = 10k) estimated by each caller for each sample (see Methods for how to compute copy number for bins from the raw CNV calls).**

**Supplementary Table 2. The data for plotting figures in Figure 3 and Supplementary Figure 2.**

**Supplementary Table 3. Purity estimates computed by the 5 computational tools at different purity levels and sequencing coverages (the data for plotting Figure 4).**

**Supplementary Table 4. Categorized copy number variation (1: amplification; 0: marginal; -1: deletion) of 287,509 bins (bin size = 10k) estimated by each caller using the median copy numbers across 21 REF replicates with a subset of bins labeled as high confidence calls.**

