## Supplemental Table 1 for "Comprehensive Assessment of Somatic Copy Number Variation Calling Using Next-Generation Sequencing Data"

**Box 1. Summary of observations and recommendations.**

| **Issue** | **Recommendation** |
| --- | --- |
| **Reference standards** | In the absence of True calls and reference sample, measuring reproducibility is an acceptable tool as a proxy for performance. We have proposed an integrative approach to establish the reference CNV Call Set that have been used in this study evaluate factors that affect the reproducibility, accuracy, and precision of CNV calling. |
| **CNV calling variability** | Variability of CNV calls is dominated by CNV callers, to increase confidence, we recommend taking consensus calls across multiple callers (e.g. n=2 or 3) and using slightly more stringent cutoffs to call amplifications (e.g. CN>3) and deletions (e.g. CN<1.5 or CN<1). However, taking consensus calls across too many callers (e.g. n>=4) or using too stringent cutoffs (e.g. CN>=4 for amplification and CN<0.5 for deletion) may greatly increase false negatives or even decrease precision (e.g. CN>7 for amplification in our analysis). We recommend that rather than using the reported CNs for downstream analysis, sorting CNV-affected regions into categories (such as amplification, deletion, and marginal) can reduce variation caused by unknown factors because the reported CNs could be unreliable especially for those regions with high CNs. |
| **Confounding factor** | Reproducibility of CNV detection is affected by experimental and technical confounding factors including impure tumor samples, FFPE-processed (versus fresh) samples, low input DNA amount, WES (versus WGS), and low sequencing coverage as well as sources of variability such as sequencing sites and CNV callers. When no confounding factor was involved, the variability of CNV calls was mainly due to callers. |
| **Prediction of Tumor Content** | Tumor purity greatly affected CNV calling by missing most calls, lowering (increasing) CNs for amplification (deletion) calls, and producing incorrect deletion calls. Low tumor purity (<= 20%) was the major confounding factor, making purity estimation unreliable |
| **FFPE vs Fresh** | FFPE-processed samples increased variability and lowered precision of CNV calls. Preparing fresh samples followed by WGS is preferred but ensuring high tumor purity is critical to have reliable CNV calls. |
| **WES vs WGS** | WES affected CNV calls by increasing variability, missing or producing incorrect calls depending on callers, and lowering the precision of deletion calls. |
| **Input DNA and Coverage** | Effects of low input DNA amount and low coverage are important but were relatively minor comparing to tumor purity and caller variation. |
| **Precision of CNV Calling** | We investigated if CNVs called by more than one caller (n>1; i.e. n=2, 3, 4, 5, or all 6 callers) would achieve higher precision and observed that precision went higher when taking consensus calls across 2 callers regardless of the influences of confounding factors. However, taking consensus calls across more callers would lower sensitivity and, thus, might lower the F1 score as precision increased. If tumor purity is less than 50%, we recommend interpreting deletions cautiously because they could be either marginal or amplification. |
| **Validation of CNV calls** | Although method evaluation studies often report the best method as their final conclusion, picking such caller for somatic CNV calling seems currently imperfect. Given the high variability of CNV calls, in addition to the approaches mentioned above, we recommend validating important CNV findings in replicates or independent datasets or gaining support from other evidences such as FISH and differential expression of affected genes. |
