## Supplementary material for "Comprehensive Assessment of Somatic Copy Number Variation Calling Using Next-Generation Sequencing Data": Table 1

Table 1. Summary of impacts of confounding factors on CNV calling^1,2,3,4^.

|  |  | **ASCAT** | **CNVkit** | **FACETS** | **FREEC** | **Segmentum** | **Sequenza** | Summary |
| --- | --- | --- | --- | --- | --- | --- | --- | --- |
| Medium tumor purity | Var+ |  | High |  | High | High |  | High |
|  | Amp |  | n.call-  CN- | CN- | n.call--  CN-- | n.call-  CN-- | n.call-  CN-- | n.call-  CN-- |
|  | Del |  | n.call-- |  | n.call--  CN+ | n.call--  CN+ |  | n.call--  CN+ |
| Low tumor purity | Var+ | Low |  |  |  | Low |  | Low |
|  | Amp | n.call---  CN--- | n.call---  CN--  P- | n.call--  CN--- | n.call---  CN--- | n.call---  CN-- | n.call--  CN-- | n.call---  CN--- |
|  | Del | n.call---  N/A  P--- | n.call---  CN++  P--- | n.call---  N/A  P--- | n.call+  CN++  P--- | n.call---  N/A  P--- | n.call-- | n.call---  CN++  P--- |
| FFPE | Var+ |  | High |  | Medium | Medium | Low | Medium |
|  | Amp |  | P- |  |  | P- | n.call-  CN-- | P- |
|  | Del | n.call+  P- |  | n.call+  P- | n.call+  P- | n.call-  P- | n.call+  P- | n.call+  P- |
| Low DNA amount | Var+ |  | Low |  |  | Low |  | Low |
|  | Amp |  |  |  |  |  | n.call-  CN-- |  |
|  | Del |  |  |  |  | n.call+  P- |  |  |
| WES | Var+ |  | Low |  | High | Medium |  | Medium |
|  | Amp | n.call++  CN+  P- |  |  | n.call--  CN-- | n.call-  CN- | n.call-  CN-- | n.call-  CN- |
|  | Del | n.call---  CN++ |  | n.call- | n.call+++  P-- | n.call+++  P-- | n.call- | n.call-/+++  P-- |
| Low coverage | Var+ |  | Medium | Low | Low | Low |  | Low |
|  | Amp |  |  | CN-- |  |  | n.call-  CN-- | CN-- |
|  | Del |  |  | n.call--  CN++ |  | n.call+++  P- |  |  |

^1^Var+ (level of variation increase compared to REF): 1 < Low <= 2 < Medium <= 5 < High (Supplementary Table 2)

^2^n.call (number of units being called relative to REF): 0 <= --- < 0.1 <= -- < 0.5 <= - < 0.8, 1.2 < + <= 1.5 < ++ <= 1.9 < +++ (Supplementary Table 2)

^3^CN (overall copy number changes for CNV calls): -1 <= --- < -0.9 <= -- < -0.5 <= - < -0.2, 0.2 < + <= 0.5 < ++ <= 0.9 <= +++ (Supplementary Table 2)

^4^P (precision): 0 <= - < 0.1 <= -- < 0.5 <= - < 0.8 (Supplementary Table 2)
