## Supplementary material for "Comprehensive Assessment of Somatic Copy Number Variation Calling Using Next-Generation Sequencing Data": Table 2

Table 2. Summary for all 6 NGS-based CNV callers.

|  | | **ASCAT** | **CNVkit** | **FACETS** | **FREEC** | **Segmentum** | **Sequenza** |
| --- | --- | --- | --- | --- | --- | --- | --- |
| Variation (REF) | | Medium | Low | Medium | Low | Low | High |
| Amplification | | 56.9% | 33.3% | 61.4% | 50.7% | 31.2% | 75.2% |
| Deletion | | 2.7% | 37.3% | 3.1% | 3.3% | 13.7% | 2.6% |
| Marginal | | 40.4% | 29.4% | 35.5% | 46% | 55.1% | 22.2% |
| ^*^F1  (M) | Amp | 0.98 | 0.8 | 0.97 | 0.96 | 0.77 | 0.87 |
|  | Del | 0.95 | 0.17 | 0.95 | 0.95 | 0.4 | 0.93 |
| ^*^F1  (A) | Amp | 0.58 | 0.84 | 0.56 | 0.64 | 0.87 | 0.45 |
|  | Del | 0.27 | 0.63 | 0.29 | 0.32 | 0.62 | 0.27 |
| Effects of confounding factors^#^ | | LP, WES | LP, MP, FFPE | LP, LC | LP, MP, WES | LP, MP, WES, LC | LP (less) |
| Additional notes for confounding factors | |  |  |  |  | Precision of deletion calls sensitive to all factors | CNs dropped for all factors |
| Purity estimation | | Not accurate at all for purity <= 20% | N/A | Low purity cannot be estimated especially for WES | N/A | N/A | Best among 5 tested tools |

^*^Median F1 when Majority calls (M) or Array calls (A) used for evaluation

^#^LP=low purity; MP=medium purity; LC=low coverage; Listed if the factor corresponds to any +++/---/high or at least 2 ++/--/medium in Table 1
